# Asymmetric Functional Gradients in the Human Subcortex

**DOI:** 10.1101/2020.09.04.283820

**Authors:** Xavier Guell, Jeremy D Schmahmann, John DE Gabrieli, Satrajit S Ghosh, Maiya R Geddes

**Author notes:** Contributed equally as senior authors (Satrajit S Ghosh, Maiya R Geddes). Correspondence: Xavier Guell.

## Abstract

A central principle in our understanding of cerebral cortical organization is that homotopic left and right areas are functionally linked to each other, and also connected with structures that share similar functions within each cerebral cortical hemisphere. Here we refer to this concept as interhemispheric functional symmetry (IHFS). While multiple studies have described the distribution and variations of IHFS in the cerebral cortex, descriptions of IHFS in the subcortex are largely absent in the neuroscientific literature. Further, the proposed anatomical basis of IHFS is centered on callosal and other commissural tracts. These commissural fibers are present in virtually all cerebral cortical areas, but almost absent in the subcortex. There is thus an important knowledge gap in our understanding of subcortical IHFS. What is the distribution and variations of subcortical IHFS, and what are the anatomical correlates and physiological implications of this important property in the subcortex? Using fMRI functional gradient analyses in a large dataset (Human Connectome Project, n=1003), here we explored IHFS in human thalamus, lenticular nucleus, cerebellar cortex, and caudate nucleus. Our detailed descriptions provide an empirical foundation upon which to build hypotheses for the anatomical and physiological basis of subcortical IHFS. Our results indicate that direct or driver cerebral cortical afferent connectivity, as opposed to indirect or modulatory cerebral cortical afferent connectivity, is associated with stronger subcortical IHFS in thalamus and lenticular nucleus. In cerebellar cortex and caudate, where there is no variability in terms of either direct vs. indirect or driver vs. modulatory cerebral cortical afferent connections, connectivity to cerebral cortical areas with stronger cerebral cortical IHFS is associated with stronger IHFS in the subcortex. These two observations support a close relationship between subcortical IHFS and connectivity between subcortex and cortex, and generate new testable hypotheses that advance our understanding of subcortical organization.

## INTRODUCTION

New methods of neuroimaging data analysis sometimes result in the observation of unexpected organizational properties in the brain. For example, (Haxby et al. 2001) showed that fusiform face area fMRI activation is sufficient to decode whether participants are presented images of either shoes or chairs.

We detected an unexpected asymmetric principal functional gradient in the human thalamus when analyzing fMRI resting-state functional connectivity using diffusion map embedding (Coifman et al. 2005). This observation was in sharp contrast with the general principle that there is high functional symmetry between left and right homotopic brain areas (Biswal et al. 1995; Lowe et al. 1998; Salvador et al. 2005; Tian et al. 2020). Here we present a succession of analyses that sought to explore the significance of this observation.

To understand why this finding was unexpected, it is necessary to describe the methodology of diffusion map embedding. Diffusion map embedding (Coifman et al. 2005) has been used recently to unmask functional gradients that represent low-dimensional macroscale properties of brain organization (Guell et al. 2018b; Margulies et al. 2016). Mathematically, these gradients represent an ordered and orthogonal set of increasingly less smooth functions on the manifold of interest (e.g. cereberal cortex, cerebellar cortex, or thalamus; for details see Shuman et al. 2013). The first gradient or “principal gradient” captures the pattern of functional connectivity that represents the smoothest gradient with the lowest amount of change, or explains the largest variance over the manifold. The second gradient will be orthogonal to the first, be less smooth, and explain lower variance than the first; and so on.

Diffusion map embedding in cerebral cortex and cerebellar cortex consistently reveals principal functional gradients that are symmetric (**Supplementary Figure 1A, B**). Both within cerebral cortex and within cerebellar cortex, these gradients progress from primary processing territories (motor areas in cerebellar cortex, sensorimotor and visual areas in cerebral cortex) to attentional and default-mode processing regions across both hemispheres (Dong et al. 2020; Guell et al. 2018b; Hong et al. 2019; Huntenburg et al. 2017; Larivière et al. 2019; Margulies et al. 2016). Gradient-based analyses of more constrained regions such as primary motor cortex (Haak et al. 2018), primary visual cortex (Haak et al. 2018), entorhinal cortex (Schröder et al. 2015), insula (Cerliani et al. 2012), temporal lobe (Bajada et al. 2017; Jackson et al. 2018), and striatum (Marquand et al. 2017) also reveal a symmetric distribution of principal functional gradients. These findings align with the general understanding that resting-state functional connectivity patterns are largely symmetric. Principal functional gradients capture the strongest progression of functional connectivity patterns within a brain structure, and so symmetry in principal functional gradients reflects the fact that there is strong symmetry in resting-state functional connectivity between left and right structures in the brain. Symmetry in fMRI resting-state functional connectivity was first discovered in motor cortex (Biswal et al. 1995). Additional studies generalized this observation to other brain regions (visual cortex, amygdala; Lowe et al. 1998), and eventually established symmetry in resting-state functional connectivity as a ubiquitous brain principle (Salvador et al. 2005). In sharp contrast to all previous gradient-based analyses of resting-state functional connectivity, an exploratory analysis in our laboratory unexpectedly revealed a principal functional gradient in thalamus that was asymmetric (**Supplementary Figure 1C**). This discovery raised the intriguing possibility that there might be strong resting-state functional connectivity asymmetries in this region of the subcortex.

What is the relevance of this observation? Functional symmetry and asymmetry in the subcortex at rest remains poorly described, and the anatomical substrate and functional significance of left-right functional symmetry in the subcortex remains poorly understood. Some studies have reported, but not discussed, subcortical patterns of resting-state asymmetry (Stark et al. 2008; Zuo et al. 2010). In contrast, many studies have characterized patterns of cerebral cortical functional symmetry and asymmetry, and analyzed the anatomical substrate and functional significance of this fundamental property of brain organization. There is more homotopic synchrony in primary sensory-motor cortices than in prefrontal and temporoparietal heteromodal association areas (Damoiseaux et al. 2006; Stark et al. 2008; Zuo et al. 2010). A large volume of evidence highlights the relevance of corpus callosum for cerebral cortical left-right functional symmetry at rest. Tracing investigations revealed that most callosal fibers connect homotopic regions of the cerebral cortex (Innocenti 1986; Pandya et al. 1971). Transection of corpus callosum in the cat and rodent abolishes cerebral cortical left-right synchrony (Engel et al. 1991; Zhou et al. 2014). Acallosal or callosotomized patients exhibit decreased interhemispheric coherence in EEG and fMRI recordings (Johnston et al. 2008; Koeda et al. 1995; Montplaisir et al. 1990; Nielsen et al. 1993; Quigley et al. 2003), and callosal lesions in multiple sclerosis result in similar consequences (Lowe et al. 2008). Additional commissural fibers in cerebral cortex such as anterior commissure (Schmahmann & Pandya 2009a) and hippocampal commissures (Schmahmann & Pandya 2009b) are likely to play a similar role. Importantly, the proposed anatomical basis of interhemispheric symmetry in the cerebral cortex is incompatible with subcortical anatomy. There is no anatomical equivalent of the corpus callosum in the subcortex - there are virtually no subcortical homotopic commissural pathways in the subcortex, with some minor exceptions such as interthalamic adhesion (Damle et al. 2017), habenular commissure (Hikosaka et al. 2008), vestibular commissures (Buttner-Ennever 1992), and commissure of the inferior colliculus (Malmierca et al. 2009). As a consequence, subcortical functional symmetry must emerge from a different anatomical reality. The functional significance of functional symmetry may also be different in the subcortex compared to cerebral cortex. In cerebral cortex, some argue that the “*high degree of synchrony observed across primary cortices reflects networks engaged in interhemispheric relay of information essential for bilateral sensory integration and motor coordination*”, while lower functional homotopy in heteromodal association areas reflects “*the predisposition of higher-order homotopic regions to operate more independently*” (Stark et al. 2008). Restated, lower and higher levels of symmetry in resting-state connectivity might correspond to lower and higher levels of functional lateralization in cerebral cortex. At the same time, interhemispheric callosal projections linking primary sensory-motor areas are more myelinated than those linking heteromodal association areas (Aboitiz et al. 1992; Lamantia & Rakic 1990), and there is stronger callosal connectivity (as indexed by diffusion-weighted MRI) in territories with lower functional lateralization as measured by fMRI task activation (Karolis et al. 2019). Left-right synchrony of a given cerebral cortical area may thus be determined not only by its functional relationship to its homotopic partner, but also by the structural features of the callosal fibers that link them together. Such an understanding of the anatomy and physiology of interhemispheric symmetry is missing in subcortical structures, as there is no detailed characterization of subcortical left-right functional symmetry, and current anatomical theories of functional symmetry are based on cerebral cortical commissural systems that are largely absent in subcortex.

Guided by the observation of an asymmetric principal functional gradient in thalamus, we set out to examine interhemispheric functional symmetry (IHFS) and interhemispheric functional asymmetry (IHFaS) in multiple subcortical brain compartments (thalamus, cerebellar cortex, caudate nuclei, and lenticular nuclei), as indexed by resting-state functional connectivity, using multiple complementary techniques (functional gradients, seed-based resting-state connectivity, and laterality indices). These measurements are influenced by multiple functional properties that together define the broad concepts of IHFS and IHFaS; this includes the fact that two homotopic left and right brain areas may be functionally linked to similar or distinct regions within each structure (as captured by functional gradients that were calculated within each subcortical structure), the related phenomenon that two homotopic left and right brain areas might be functionally linked preferentially to one hemisphere (as captured by laterality indices calculated within each subcortical structure), and the closely linked phenomenon that two homotopic left and right brain areas may be strongly or weakly connected to each other. Asymmetry in functional gradients served as an anchor to guide these analyses. These techniques can characterize IHFS/IHFaS variations across these structures, and address the knowledge gap that functional symmetry variations across subcortex are not well characterized. New knowledge emerging from this characterization may be used to build novel theories of the anatomical and physiological significance of IHFS in the subcortex.

## METHODS

### Functional gradients

Functional gradient analyses were performed on the data provided by the Human Connectome Project (HCP), WU-Minn Consortium (Van Essen et al. 2013), including 1003 healthy participants. Functional gradient calculation from resting-state fMRI data is an analytical strategy based on novel dimensionality reduction techniques (Coifman et al. 2005) that has unmasked fundamental properties of brain macro-scale organization in studies of cerebral cortex (Margulies et al. 2016) and cerebellum (Guell et al. 2018b). This methodology analyzes the similarity between the connectivity patterns of each datapoint within a given brain structure, and extracts functional gradients that represent the principal poles and transitions of connectivity patterns within that structure. Our analysis started by restricting the whole-brain group-average resting-state connectivity matrix provided by HCP to each of the structures of interest: cerebellar cortex, caudate nuclei, thalami, and lenticular nuclei (comprising the putamen and globus pallidus externa and globus pallidus interna). We thus obtained four connectivity matrices, one for each of these four structures, which included functional connectivity values from each voxel to all other voxels within that structure. Within each of these structures, the “connectivity pattern” of each voxel was represented as an n-dimensional vector, where n corresponded to the total number of voxels in that structure. Because all voxels within each structure were represented in the same n-dimensional space, cosine distance between each pair of vectors could be calculated, and an affinity matrix was constructed as (1 – cosine distance) for each pair of vectors. This affinity matrix represented the similarity of connectivity patterns for each pair of voxels within each structure. A Markov transition matrix was then constructed using information from the affinity matrix; information from the affinity matrix was thus used to represent the probability of transition between each pair of vectors within each structure. In this way, there was a higher transition probability between pairs of voxels with similar connectivity patterns. This probability of transition between each pair of voxels was analyzed as a symmetric transformation matrix, allowing the calculation of eigenvectors on the Laplacian of this matrix. Eigenvectors derived from this transformation matrix represented the principal orthogonal directions of transition between all pairs of voxels within each structure. See https://github.com/satra/mapalign for an implementation of this methodology, and our online repository for the application of this methodology to our data (https://github.com/xaviergp/subcortical_IHFS).

Functional gradient values indicate the similarity of functional connectivity patterns of all datapoints (voxels or vertices) within a given brain structure. Two brain coordinates with similar functional gradient values will have similar functional connectivity patterns. In this way, functional gradients can identify regions of high IHFaS if the distribution of functional gradient values is asymmetric in left versus right homotopic areas. Conversely, functional gradients can identify regions of high IHFS if functional gradient values are similar in left versus right homotopic areas. Functional gradient analysis identifies a first component (or “principal functional gradient”) that accounts for as much of the variability in the data as possible. Each following component (each following gradient) accounts for the highest variability possible under the constraint that all gradients are orthogonal to each other. Our analyses were focused on the principal functional gradient of each structure, as this component accounts for the highest amount of variability and is calculated first with no orthogonality constraints. Non-principal functional gradients were explored in additional analyses. Contrasting with other measures of homotopic synchrony such as Voxel-Mirrored Homotopic Connectivity (Zuo et al. 2010), functional gradient calculations do not require the transformation of brain data into a symmetric brain template. Results within thalamus were mapped using an MRI atlas of thalamic nuclei (Krauth et al. 2010).

### Laterality index and seed-based functional connectivity maps

To further examine the relationship between functional gradients and IHFS, seed-based functional connectivity maps were computed from each voxel within each structure (thalami, lenticular nuclei, caudate nuclei, cerebellar cortex). These maps were used to calculate a laterality index as follows: (left_score – right_score) / (left_score + right_score), where left_score and right_score correspond to the sum of all functional connectivity values for each left and right structure (for example, in the case of thalamus, functional connectivity values in left and right thalamus). The relationship between functional gradients and laterality indices were analyzed using scatterplots. To provide an exemplar visualization of IHFS differences between distinct regions of a structure (e.g., between distinct regions of thalamus), a small number of seed-based functional connectivity maps were visualized from specific datapoints within that structure. These datapoints were selected based on functional gradient values as explained in the Results section. Unthresholded z correlation maps from these datapoints were generated and visualized using the workbench_view graphical user interface (Marcus et al. 2013). Note that thresholds based on statistical significance (such as p values) are not useful in the context of a sample of 1003 participants; because of the very large sample size, even very small effects would be statistically significant.

### Projection of subcortical functional gradients to cerebral cortex

To allow the comparison of asymmetric functional gradients from different subcortical structures in a common anatomical space, and to examine the relationship between these functional gradients and cerebral cortical functional connectivity, subcortical functional gradients were projected to cerebral cortex, as follows. For each subcortical functional gradient (G) and each subcortical voxel (V), the cerebral cortical functional connectivity map of each V was multiplied by the absolute G value of V. For example, for cerebellar cortical functional gradient, cerebral cortical functional connectivity map of voxel A was multiplied by the absolute functional gradient value of voxel A; cerebral cortical functional connectivity map of voxel B was multiplied by the absolute functional gradient value of voxel B; and so on for each voxel in cerebellar cortex. Then, all weighted cerebral cortical functional connectivity maps (as many as the total number of voxels in cerebellar cortex) were added together. The resulting cerebral cortical maps provided a visualization of the significance of asymmetric subcortical functional gradients in terms of functional connectivity to cerebral cortex. Absolute functional gradient values were used in order to specifically observe the relationship between subcortical regions with strong IHFaS as indexed by asymmetric functional gradients and cerebral cortical connectivity.

### Data and code access

All data used to perform analyses in the present study are publicly available as part of the HCP dataset, facilitating future replications or re-analyses of our findings. Thalamic nuclear MRI atlas can be obtained by contacting the authors of the atlas (Krauth et al. 2010). Custom code, intermediate files, and final functional gradient and seed-based connectivity maps from our analyses are openly available at https://github.com/xaviergp/subcortical_IHFS.

## RESULTS

### Principal functional gradients are distributed symmetrically in cerebellar cortex and caudate; and distributed asymmetrically in certain homotopic regions of thalamus and lenticular nucleus

The principal functional gradient was examined first within each structure (cerebellar cortex, caudate nuclei, thalami, lenticular nuclei). The principal functional gradient captures the highest possible amount of data variability, and is the only component that is free from orthogonality constraints because it is the first functional gradient to be calculated (all functional gradients calculated afterwards – i.e. functional gradients 2, 3, etc. – must be orthogonal to previous functional gradients).

As expected from previous studies (Guell et al. 2018b), the principal functional gradient was distributed in a largely symmetric fashion in all territories of cerebellar cortex (**Figure 1**). Principal functional gradient in caudate nuclei was also distributed symmetrically in left and right caudate, and followed a rostral-to-caudal progression (**Figure 1**). In sharp contrast, distribution of the principal functional gradient was asymmetric in thalami and lenticular nuclei. Importantly, regions of asymmetry in thalami and lenticular nuclei were homotopic (i.e. located in the same territory in the right and left hemisphere); regions of left thalamus that contained the lowest functional gradient values (green color in **Figure 1**) were homotopic to the regions of right thalamus that contained the highest functional gradient values (yellow color in **Figure 1**). The same was true for lenticular nucleus.

**Figure 1.**
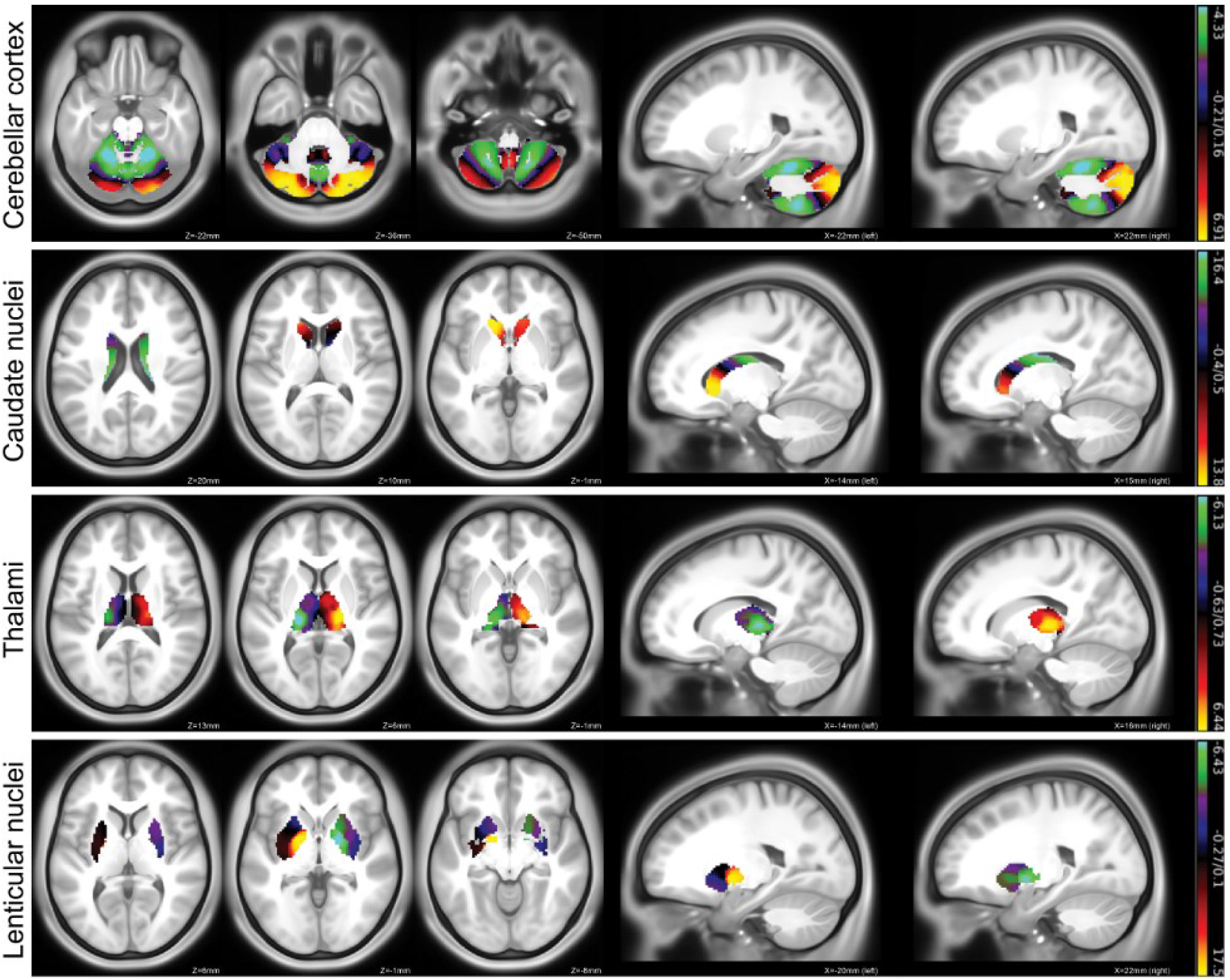
<u>Distribution of functional gradients in cerebellar cortex, caudate nuclei, thalami, and lenticular nuclei</u>. A symmetric distribution is observed in cerebellar cortex, and caudate nuclei – note that the distribution of color intensity is largely symmetric in these cases; higher (yellow color) and lower (green color) poles of functional gradient values are present in both right and left components of these structures. In contrast, functional gradient distribution is asymmetric in thalamus and lenticular nucleus – in these two cases, each pole of the distribution of functional gradient values (yellow color and green color) is located exclusively in either the right or the left hemisphere. Importantly, regions of asymmetry in thalami and lenticular nuclei are homotopic – namely, the region of left thalamus that contained the lowest functional gradient values (green color) is homologous to the regions of right thalamus that contained the highest functional gradient values (yellow color), and the same is true for lenticular nucleus. See Supplementary Figure 2 for a detailed analysis of gradient values within each thalamic nucleus.

Within lenticular nuclei, regions of asymmetry corresponded to pallidum, but not to putamen (**Figure 1**). Within the thalami, regions of asymmetry corresponded mostly to first order nuclei including ventral posterior nuclei, but not to higher order nuclei such as medial dorsal and anterior thalamic nuclei (a detailed description of first order vs. higher order thalamic nuclei is provided in the discussion section). While this pattern within thalamus is observable in **Figure 1**, a more detailed quantification of functional gradient values in each thalamic nucleus using an MRI atlas (Krauth et al. 2010) is shown in **Supplementary Figure 2** (for reference regarding thalamic anatomy, see Felten et al. 2016, Guillery 1995).

### Asymmetric functional gradients in thalamus and lenticular nucleus reflect variations in interhemispheric symmetry within these structures as measured by laterality index

To further characterize the relationship between principal functional gradients in thalami and lenticular nuclei and IHFS/IHFaS, we computed seed-based functional connectivity maps from each voxel within thalami and lenticular nuclei. We then computed a laterality index that quantified the degree of asymmetry in each functional connectivity map from each seed (see methods), and plotted laterality index scores for each voxel in thalami and lenticular nuclei against their corresponding functional gradient value. This analysis showed that more extreme positions in the principal functional gradient of thalami and lenticular nuclei correspond to areas where there is stronger asymmetry of functional connectivity (**Figure 2**).

**Figure 2.**
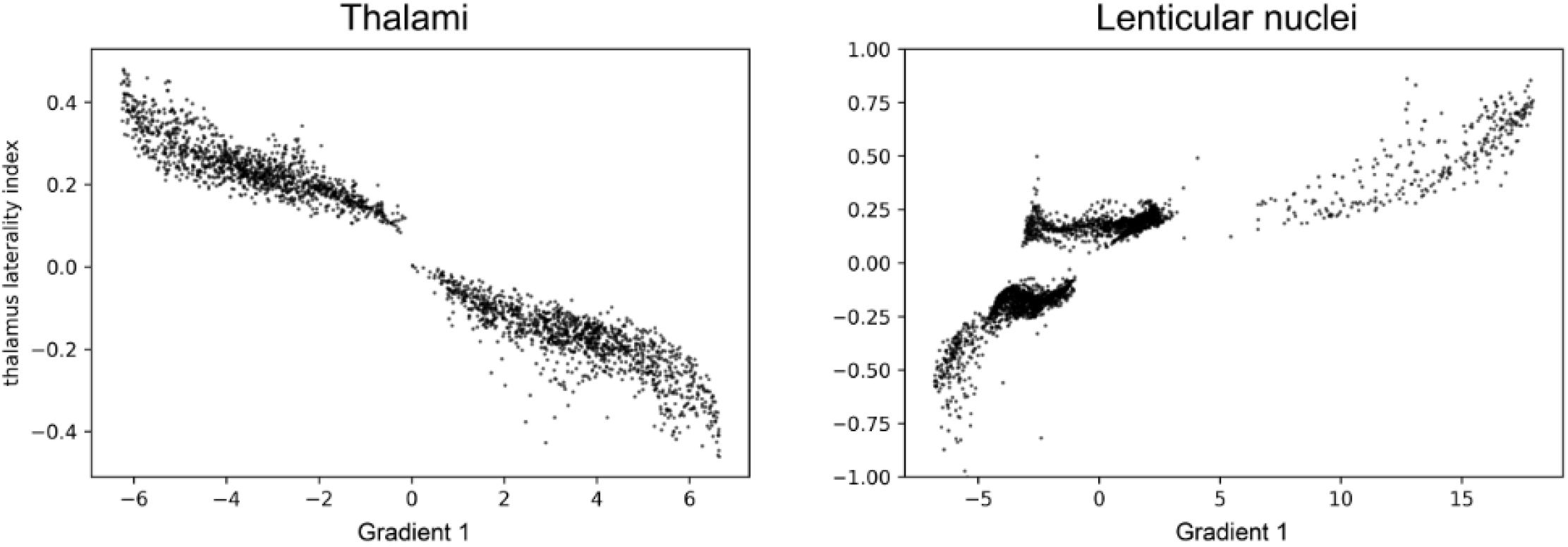
<u>Laterality index compared to principal functional gradient in thalami and lenticular nuclei</u>. More extreme positions in the principal functional gradient of thalami and lenticular nuclei correspond to areas where there is stronger lateralization of functional connectivity.

To obtain exemplar illustrations of the finding reported in **Figure 2**, we visualized functional connectivity maps calculated from the highest and lowest functional gradient values in thalamus (MNI x, y, z = 20, -22, 2; -18, -26, 4) and lenticular nucleus (16, -2, -2; -14, -6, -4). As expected, functional connectivity maps from these seeds were asymmetric (**Figure 3**, first and third rows). These maps were compared to functional connectivity maps calculated from regions of thalamus and lenticular nucleus that possessed medium (as opposed to extreme) functional gradient values. Thalamic and lenticular regions of medium functional gradient values were identified as follows. Lowest values of thalamic principal functional gradient were located in left thalamus, and thus medium functional gradient values in left thalamus were identified as the highest principal functional gradient value within left thalamus (which corresponded to MNI x, y, z = -2, -12, 12). Highest values of thalamic principal functional gradient were located in right thalamus, and thus medium principal functional gradient values in right thalamus were identified as the lowest principal functional gradient value within right thalamus (14, -30, -4). The same strategy was used in lenticular nucleus (left = -28, 4, -4; right = 30, -8, -10). As expected, functional connectivity maps from regions of thalamus and lenticular nucleus that possessed medium (as opposed to extreme) functional gradient values were comparatively less asymmetric (**Figure 3**, second and fourth rows).

**Figure 3.**
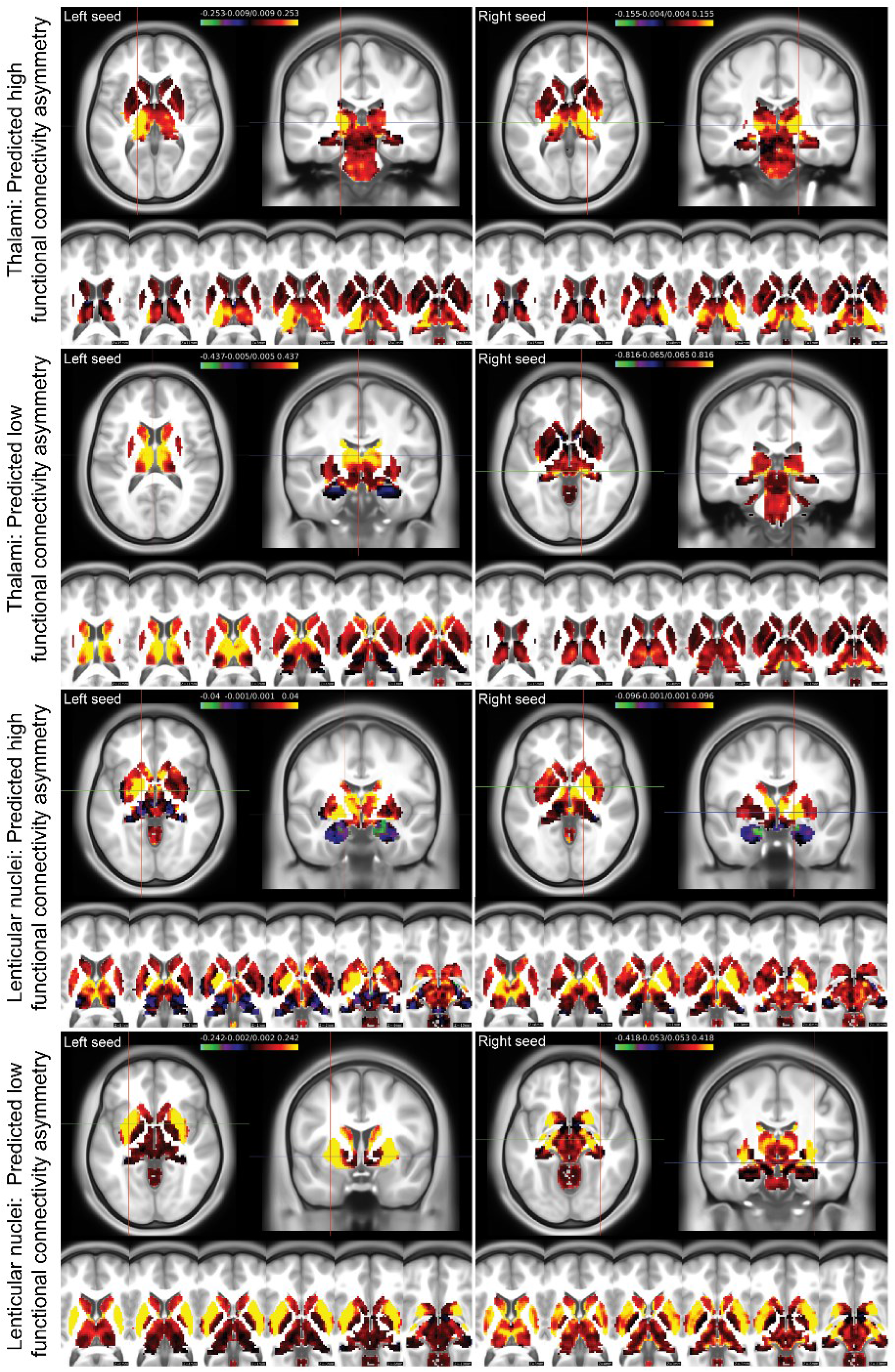
<u>Functional connectivity calculated from principal functional gradient extreme (maximum/minimum) values (predicted high functional connectivity asymmetry) and medium values (predicted low functional connectivity asymmetry) in thalami and lenticular nuclei</u>. Seed region for each functional connectivity map is indicated by a crosshair in the axial and coronal view located in the upper half of each panel. Additional axial cuts are shown for each connectivity map in the lower half of each panel. As predicted, distribution of functional connectivity strength maps indicate that principal functional gradient maximum/minimum values correspond to areas with higher functional connectivity asymmetry, while principal functional gradient medium values correspond to areas with lower functional connectivity asymmetry. All subcortical brain territories are included in this figure, according to the HCP definition of subcortical structures. First row left: functional connectivity strength is superior in left thalamus than in homotopic aspects of right thalamus when seeding from the coordinate of predicted low functional connectivity symmetry. First row right: while some strong functional connectivity (yellow color) is observed in the contralateral left homotopic thalamic region, functional connectivity map is asymmetric with more functional connectivity in right thalamus (see axial cuts in the lower half of the image). Second row: connectivity maps are symmetric for left and right thalamic areas of predicted low functional connectivity asymmetry. Third and fourth rows: as in thalamus, connectivity maps from lenticular coordinates of predicted high functional connectivity asymmetry (third row) are more asymmetric than connectivity maps from lenticular coordinates of predicted low functional connectivity asymmetry (fourth row).

Functional connectivity maps shown in **Figure 3** provide an alternative visualization of the same result that is reported in **Figure 2**, but note that **Figure 3** includes only functional connectivity calculated from four voxels in thalamus and lenticular nucleus, while **Figure 2** includes all voxels in thalamus and lenticular nucleus.

### Non-principal functional gradients in cerebellum (third gradient) and caudate (second gradient) are distributed asymmetrically, and reflect variations in IHFS as measured by laterality index

Following an analysis of the principal functional gradients of each structure, we explored whether non-principal functional gradients (namely, functional gradients calculated after the principal gradient, which must be orthogonal to previous functional gradients) exhibited an asymmetric pattern. As expected from our previous report (see supplementary material in Guell et al. 2018b), the third functional gradient in cerebellum exhibited an asymmetric pattern. An asymmetric pattern was also observed in the second functional gradient in caudate (**Figure 4**). When analyzing the relationship between laterality indices and these asymmetric functional gradients in cerebellum and caudate, a similar relationship as described in **Figure 2** was observed, in which extreme values of the asymmetric gradients corresponded to areas with more lateralized functional connectivity maps (**Figure 4D**; these relationships were not as clear as in **Figure 2**, but were nonetheless appreciable).

**Figure 4.**
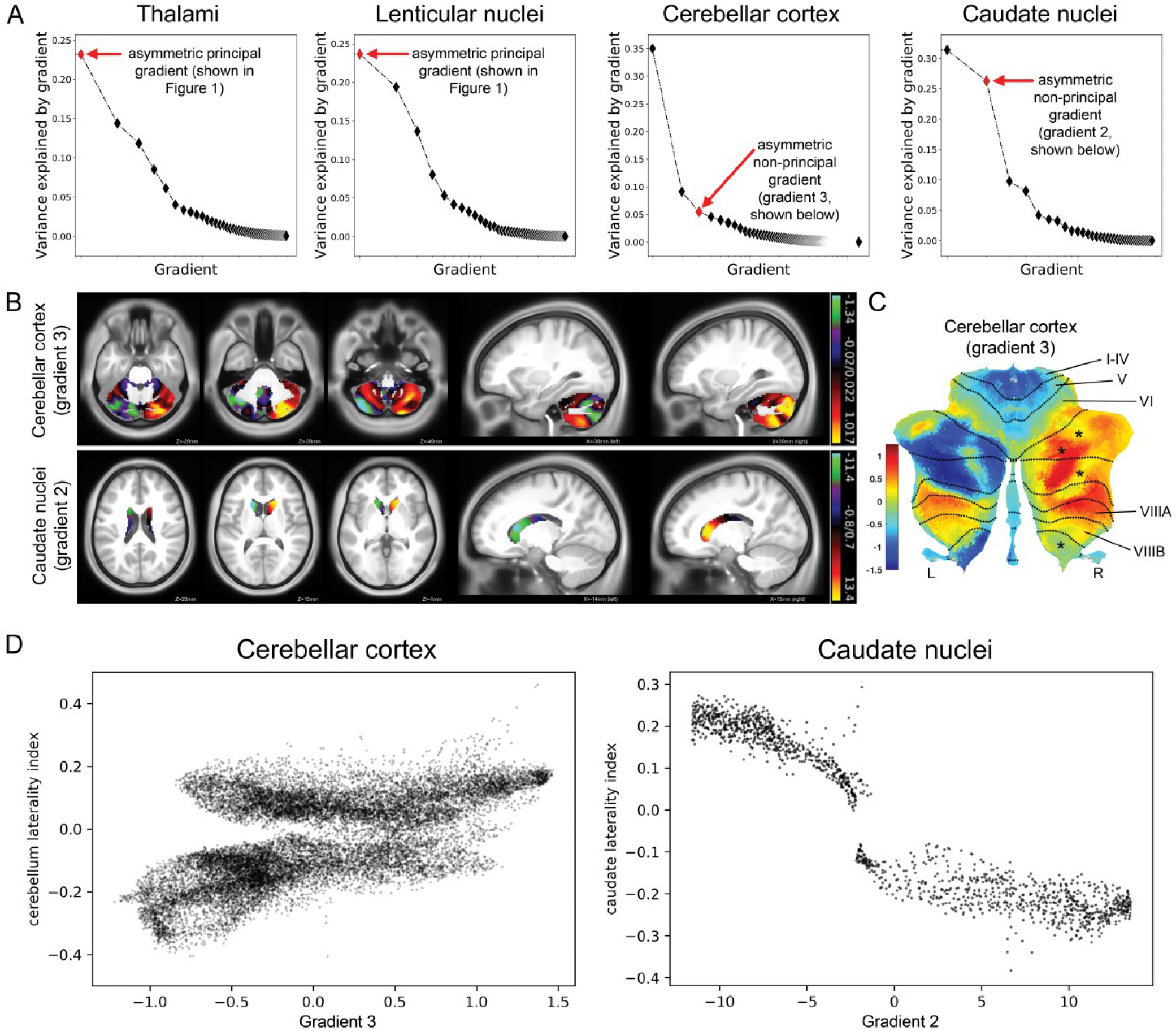
<u>Non-principal functional gradients are distributed asymmetrically in cerebellar cortex (third gradient) and caudate nuclei (second gradient)</u>. **(A)** Variance explained by each functional gradient; red diamonds correspond to the earliest asymmetric functional gradient. The principal (i.e. first) functional gradient was asymmetric in thalami and lenticular nuclei, as shown in Figure 1. When visualizing non-principal functional gradients (i.e. functional gradients 2, 3, etc.), the earliest functional gradient to reveal an asymmetric distribution in cerebellar cortex and caudate nuclei were functional gradients 3 and 2, respectively. **(B)** Functional gradient 3 in cerebellar cortex reveals an asymmetric distribution; specifically, some homotopic regions of cerebellar cortex are located at opposite ends of the spectrum of functional gradient 3 (red/yellow versus purple/green). The same is true in functional gradient 2 of caudate nuclei. **(C)** Plotting cerebellar gradient 3 results in a flatmap (Diedrichsen & Zotow 2015) provides a better visualization of cerebellar gradient 3 distribution. Specifically, gradient 3 is symmetric predominantly in territories of cerebellar motor control (namely lobules I-VI and VIII (Guell et al. 2018a), indicated in the figure). The rest of territories in cerebellar cortex correspond to non-motor control, and in these territories gradient 3 is predominantly asymmetric (note how territories in cerebellar right hemisphere marked with an asterisk show different functional gradient 3 values in their corresponding homotopic left hemisphere regions). **(D)** Laterality index plotted in relation to asymmetric functional gradients in cerebellar cortex and caudate nuclei. More extreme positions in the asymmetric functional gradient of cerebellar cortex and caudate nuclei correspond to areas where there is stronger lateralization of functional connectivity; these relationships are not as clear as in Figure 2, but nonetheless remain observable.

### Subcortical asymmetric gradients projected to cerebral cortex reveal that cerebellar and caudate areas with high IHFaS, but not lenticular and thalamic areas with high IHFaS, project to cerebral cortical areas with high IHFaS

Projecting subcortical asymmetric gradients to cerebral cortex revealed that regions with IHFaS in cerebellum and caudate map predominantly to default-mode and frontoparietal networks of the cerebral cortex (**Figure 5A**). In contrast, regions with IHFaS in thalamus and lenticular nucleus mapped predominantly to ventral attention network (**Figure 5A**). Previous studies indicate that frontoparietal and default-mode network in cerebral cortex exhibits higher IHFaS when compared to other aspects of the cerebral cortex, including attention networks (Stark et al. 2008; Zuo et al. 2010). The same dissociation remained observable when quantifying laterality index values in cerebral cortex using a threshold of 90th percentile for projected subcortical gradients (i.e. calculating laterality index in those cerebral cortical areas that were included within the boundaries of the 90th percentile of each subcortical gradient projected to cerebral cortex) (**Figure 5B**). The principal functional gradient of cerebral cortex captures a progression from motor to default-mode network (Margulies et al. 2016). A relationship between laterality indices and cerebral cortical principal functional gradient is thus expected. This expectation was confirmed in our data (**Figure 5C**). Given this close relationship between laterality indices and cerebral cortical principal functional gradient, we were also able to observe that regions of IHFaS in cerebellum and caudate, but not of thalamus and lenticular nucleus, map to cerebral cortical areas of high cerebral cortical principal gradient values (i.e. regions that are at or close to default-mode network) (**Figure 5D**).

**Figure 5.**
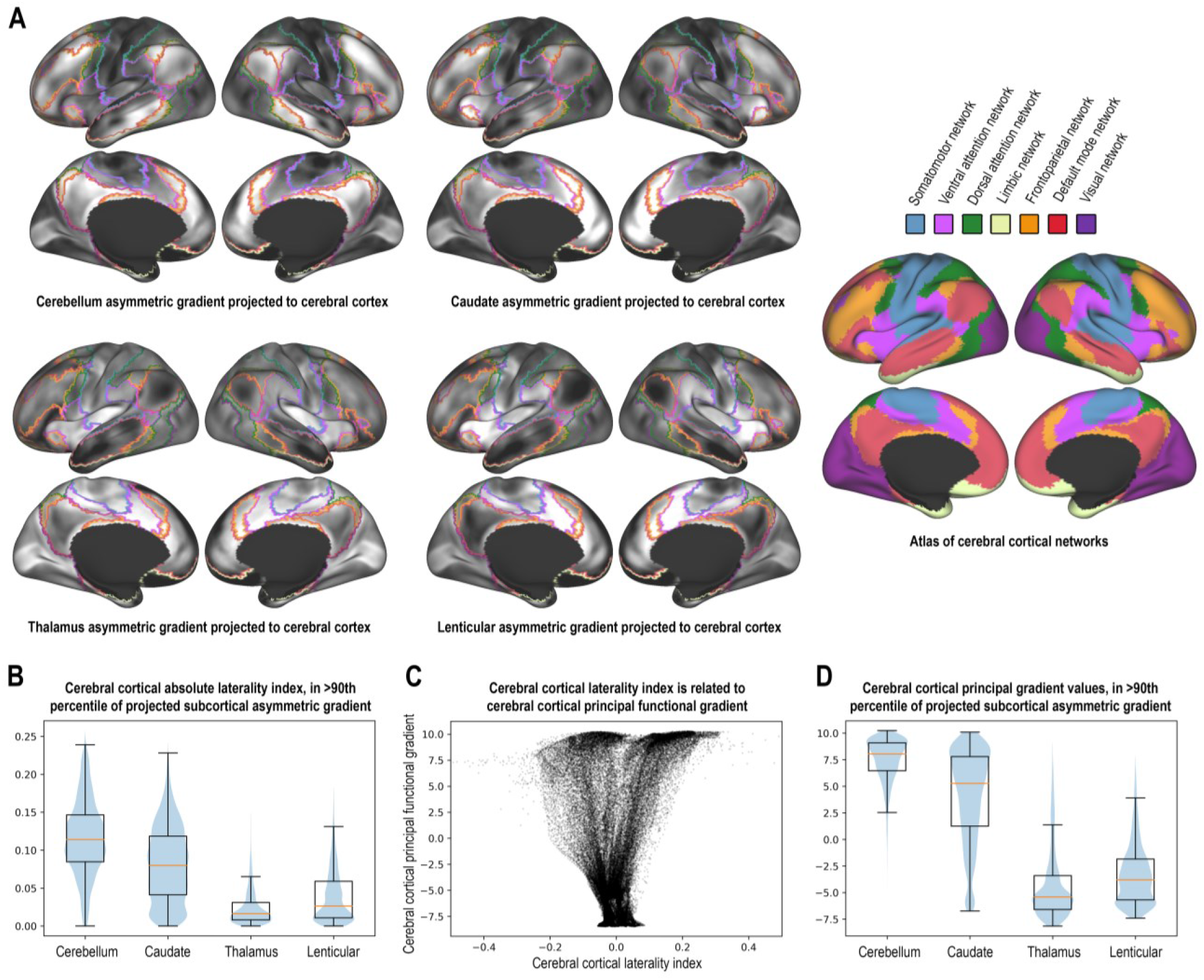
<u>Projecting subcortical asymmetric gradients to cerebral cortex</u>. **(A)** White pole of black-to-white color scale corresponds to regions of cerebral cortex that reveal stronger functional connectivity to subcortical regions of higher IHFaS (as indexed by asymmetric functional gradients in the subcortex). Top two rows: regions of IHFaS in cerebellum and caudate map predominantly to frontoparietal and default-mode networks of the cerebral cortex (red and orange borders). Bottom two rows: regions of IHFaS in thalamus and lenticular nucleus map to other cerebral cortex areas (predominantly ventral attention network; light purple border). A cerebral cortical network atlas is shown in the third column (from Yeo et al. 2011); note that borders of each network are shown in first and second columns, and that colors of each border correspond to the colors of the atlas shown in the third column. **(B)** When quantifying cerebral cortical laterality index values present within the boundaries of the top 90% percentile of projected gradients in cerebral cortex, cerebellum and caudate but not lenticular and thalamic regions of IHFaS demonstrated a preference towards cerebral cortical areas of higher IHFaS as indexed by cerebral cortical laterality indices. **(C)** Larger cerebral cortical principal functional gradient values (i.e. regions closer to default-mode network) are associated with higher laterality indices (either positive or negative; positive indicates preference towards left hemisphere and negative indicates preference towards right hemisphere). This observation is consistent with prior literature (Stark et al. 2008; Zuo et al. 2010) showing higher interhemispheric asynchrony in frontoparietal and default mode areas, since higher principal functional gradient values in cerebral cortex correspond to regions close to or at frontoparietal or default-mode network (Margulies et al. 2016). Note a shift of the laterality index values towards positive values (i.e. towards left hemisphere) at the extreme positive pole of the cerebral cortical principal functional gradient (i.e. default mode network areas); this observation is consistent with prior literature reporting a left-lateralization of DMN in cerebral cortex (Agcaoglu et al. 2014). **(D)** A dissociation similar to (B) was observed when analyzing cerebral cortical principal gradient values: cerebellum and caudate but not thalamus and lenticular projected to regions of higher cerebral cortical principal gradient values; this relationship is expected given the associations reported in (B) and (C).

## DISCUSSION

Subcortical interhemispheric functional symmetry (IHFS) and asymmetry (IHFaS) remain poorly described. The proposed anatomical basis of these properties that is based on cerebral cortical callosal and other commissural connectivity are incompatible with subcortical anatomy. Here we present a detailed analysis of IHFS/IHFaS in thalamus, lenticular nucleus, cerebellar cortex, and caudate nucleus. The synopsis of the findings discussed below is that direct (monosynaptic, as opposed to indirect/polysynaptic) and driver (as opposed to modulatory) afferent anatomical connectivity from the cerebral cortex is associated with stronger subcortical IHFS in thalamus and lenticular nucleus. In the cerebellar cortex and caudate nucleus that have no variability in terms of either direct vs. indirect or driver vs. modulatory cerebral cortical afferent connections, stronger subcortical IHFS is associated with connectivity to cerebral cortical areas that themselves have stronger IHFS. Taken together, these observations support a strong relationship between subcortical IHFS/IHFaS and connectivity between subcortex and cortex, and generate new testable hypotheses that advance our understanding of subcortical organization. A summary of our observations and interpretations is provided in **Table 1**.

**Table 1.** Summary of our observations and corresponding interpretations in thalamus, lenticular nucleus, cerebellar cortex, and caudate. IHFS = interhemispheric functional symmetry. IHFaS = interhemispheric functional asymmetry.

|  | Observation | Relevant background | Interpretation (i) | Interpretation (ii) |
| --- | --- | --- | --- | --- |
| Thalamus | IHFaS is more prominent in first order nuclei (compared to higher-order nuclei) (Figs. 1, 2, 3, Supp Fig. 2). | First order nuclei receive cerebral cortical modulatory afferents but not cerebral cortical driver afferents. Higher order nuclei receive both modulatory and driver cerebral cortical afferents. | In structures that have **variability** in terms of either direct vs. indirect or driver vs. modulatory cerebral cortical afferents (thalamus and lenticular nucleus), IHFaS is stronger in regions that have no direct or driver cerebral cortical input. | There is a strong relationship between subcortical IHFS/IHFaS and connectivity between subcortex and cortex. |
| Lenticular nucleus | IHFaS is more prominent in pallidum (compared to putamen) (Figs. 1, 2, 3). | Pallidum does not receive direct (monosynaptic) cerebral cortical anatomical projections, whereas the putamen does. |  |  |
| Cerebellar cortex | IHFaS is more prominent in territories linked to cerebral cortical areas with higher IHFaS (Figs. 4, 5); this is true in both cerebellar cortex and caudate. | In cerebellar cortex and caudate there is no variability in terms of direct (monosynaptic) vs. indirect (polysynaptic) cerebral cortical afferents, and also no variability in terms of driver vs. modulatory cerebral cortical afferents. | In structures that have **no variability** in terms of either direct vs. indirect or driver vs. modulatory cerebral cortical afferents (cerebellar cortex and caudate nucleus), IHFaS is stronger in regions that are linked to cerebral cortical areas that have higher IHFaS. |  |
| Caudate |  |  |  |  |

### In thalami and lenticular nuclei, IHFaS is stronger in areas without direct or driver afferent anatomical connections from cerebral cortex

Our analyses revealed strong IHFaS in pallidum but not in putamen, and in first order thalamic nuclei but not in higher order thalamic nuclei (**Figures 1, 2, Supplementary Figure 2**). Previous studies have shown a similar dissociation, but they were focused on cerebral cortical IHFS and did not address subcortical patterns. Stark and colleagues calculated functional connectivity between 112 regions in cerebral cortex and subcortex; their findings situated pallidum as one of the brain regions with the lowest degree of homotopic interhemispheric correlations (see third figure in (Stark et al. 2008)). In that study, all voxels within thalamus were averaged together. Zuo and colleagues calculated whole-brain voxel-mirrored homotopic connectivity maps, and provided figures where it was readily apparent that pallidum and some caudal aspects of the thalami (grossly matching thalamic territories identified in our analysis) exhibited low homotopic synchrony when compared to putamen and other aspects of thalamus (see first figure, volumetric slices in the left panel in (Zuo et al. 2010)). One recent study analyzed functional gradients in the subcortex but did not identify asymmetric functional gradients (Tian et al. 2020), likely due to the fact that many subcortical structures were grouped together in a single analysis space (rather than analyzed individually, as in this study). Another examined functional gradients in thalamus and also did not identify asymmetric functional gradients (Yang et al. 2020). We suspect this was because gradients were calculated using functional connectivity data from thalamus to cerebral cortex. Our approach differed in that we performed gradient analyses using functional connectivity data within each structure of interest (e.g. functional connectivity within thalamus, within cerebellum, etc.).

The observation of strong IHFaS in two different aspects of two distinct subcortical structures (thalamus and lenticular nucleus) provided an opportunity to interrogate the anatomical and physiological basis of IHFaS in the subcortex. We set out to identify anatomical or physiological similarities or differences that capture the relationship of the pallidum to putamen on the one hand, and the first order to higher order thalamic nuclei on the other. Our findings indicate that the feature that distinguishes the pallidum - putamen relationship and the first order - higher order thalamic relationship, and that is common to both, is the nature of their cerebral cortical inputs, specifically, the distinction between direct vs. indirect cerebral cortical afferents in lenticular nucleus, and the distinction between driver vs. modulatory cerebral cortical afferents in thalamus.

The concept of driver vs. modulatory cerebral cortical afferents was introduced originally with respect to input to thalamus (Guillery 1995) and may be explained as follows. First order (sensory) thalamic nuclei are the sensory nuclei that receive their afferents from the periphery, that is, the external world. The medial lemniscus carries tactile and proprioception information to the ventral posterior nuclei, the retina conveys visual information to the lateral geniculate nucleus, and the inferior colliculus provides auditory information to the medial geniculate nucleus. These peripheral inputs to the first order thalamic nuclei are regarded as driver afferents. The sensory thalamic nuclei are also engaged in heavy reciprocal interactions with the cerebral cortex, but the functional properties of the corticothalamic afferents to these first order thalamic nuclei are regarded not as driver (which arise in the periphery), but rather modulatory. In contrast to the first order thalamic nuclei, the higher order (associative) thalamic nuclei such as the medial dorsal nucleus, anterior thalamic nuclei, lateral posterior nucleus, and pulvinar nuclei, have no input from the external world, and are engaged instead in reciprocal corticothalamic interactions with the cerebral cortical association areas (Jones 1985; Schmahmann 2003). The cortical afferents to these higher order thalamic nuclei are the equivalent to the peripheral inputs to the sensory nuclei, and are regarded as driver afferents.

There are important differences in the cytoarchitectonic areas and laminar origins of the cerebral cortical inputs to these first order / sensory vs higher order / associative thalamic nuclei. Driver fibers emerge from layer V, whereas modulatory fibers emerge from layer VI (Abramson & Chalupa 1985; Conley & Raczkowski 1990; Gilbert & Kelly 1975; Guillery 1995; Sherman 2016; Xiao et al. 2009). First order, sensory, thalamic nuclei thus receive their cortical inputs almost exclusively from layer VI, subserving a modulatory function. Higher order thalamic nuclei receive both driver inputs from layer V, and modulatory inputs from layer VI (Xiao et al., 2009).

Driver and modulatory inputs are distinguishable at other levels of investigation as well. At the histological level, driver fibers have large terminals and round synaptic vesicles with large dendritic stems close to the cell body, whereas modulatory fibers are finer and have less complex terminal structures which lie far from the cell body. The patterns of degeneration following fiber transection also differ. Driver fibers have a relatively slow set of degenerative changes, modulatory fibers degenerate rapidly. The electrophysiological properties are also specific to each type, including differences in excitatory postsynaptic potential amplitude, paired-pulse effects and probability of transmitter release (as reviewed in Guillery 1995; Sherman 2016).

Note that whereas there is variability in corticothalamic projections of driver vs modulatory input, there is no evidence of variability in terms of driver vs modulatory input in cerebral cortical afferent fibers to the cerebellum (corticopontine fibers, Glickstein et al. 1985; Schmahmann & Pandya 1991, 1995, 1997; Schmahmann et al. 2004), caudate nucleus, putamen, or pallidum (corticostriatal fibers, Levesque 1998; McFarland & Haber 2000; Yeterian & Pandya 1994). This is despite the fact that there are cytoarchitectonic variations between the different cerebral cortical areas that project to topographically organized territories within these systems (e.g. Chikama et al. 1997), that only some portions of cerebellar cortex receive input from sensorimotor and vestibular systems (Schmahmann et al. 2019), and that different functional territories are engaged in motor, cognitive, or affective behaviors within the cerebellum (Buckner et al. 2011; Guell et al. 2018a; Schmahmann & Pandya 1997a; Stoodley et al. 2012, 2016), caudate, putamen, and pallidum (Choi et al. 2012; Parent 1990).

A different kind of dissociation in the organization of cerebral cortical afferents is present in the lenticular nucleus. Here, whereas cerebral cortical projections to the putamen are direct (monosynaptic), cerebral cortical inputs to the pallidum are indirect via intervening synaptic steps (see Felten et al. 2016; Redgrave et al. 2010).

It therefore appears that there is a shared anatomical property in the relationship [first order vs. higher order thalamic nuclei] and [pallidum vs. putamen] in that the first order thalamic nuclei and pallidum lack direct or driver afferent anatomical connections from cerebral cortex.

We illustrate our interpretation of these observations as they relate to interhemispheric functional symmetry or asymmetry in **Figure 6A** and **Table 1 (top two rows**): in thalami and lenticular nuclei, lack of direct or driver afferent anatomical connections from cerebral cortex is associated with stronger IHFaS.

**Figure 6.**
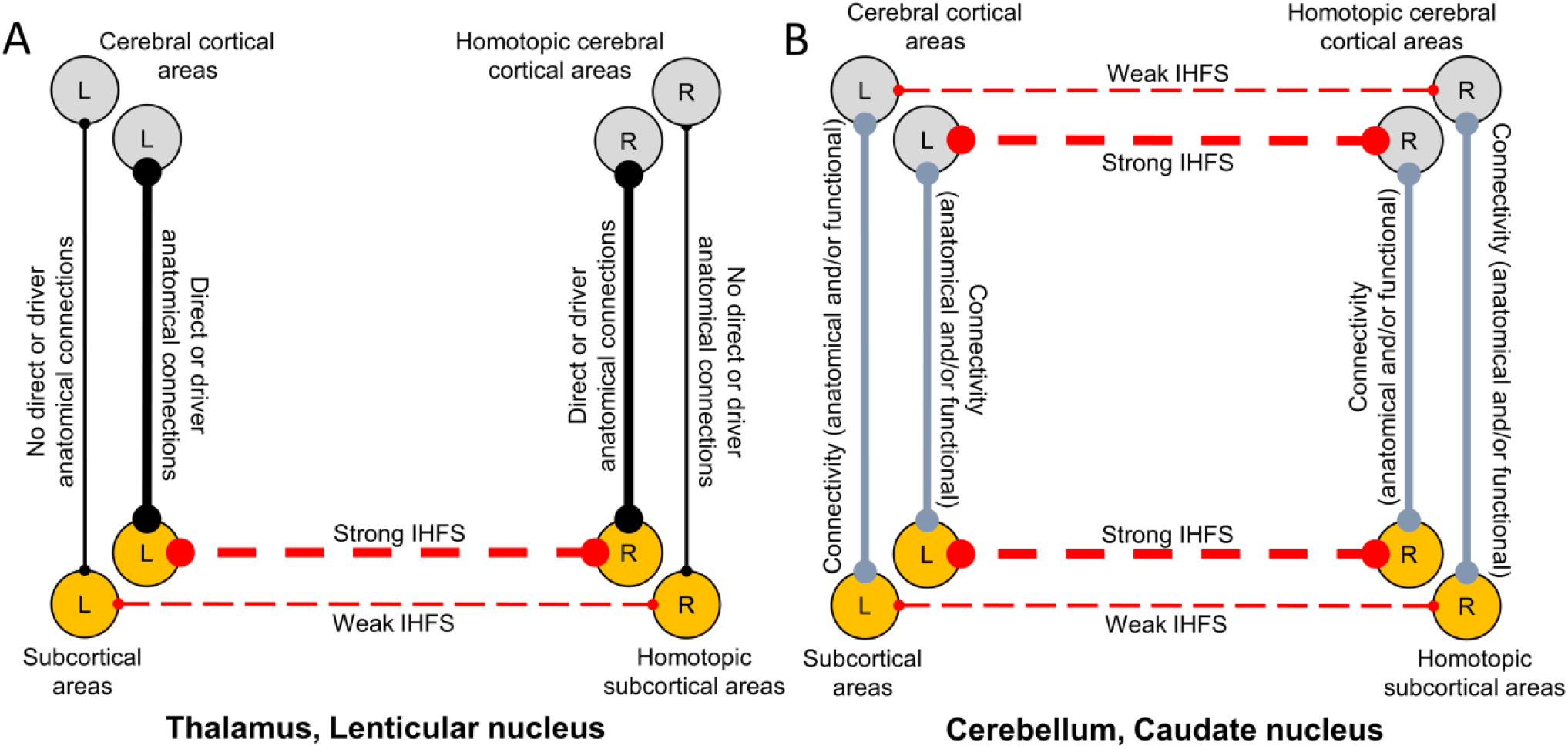
<u>Diagram of the interpretation of our observations.</u> **(A)** In subcortical territories with variability in terms of either direct vs. indirect or driver vs. modulatory cerebral cortical afferent connections, direct or driver afferent anatomical connections from cerebral cortex are associated with stronger IHFS. This conclusion is based on our results in thalami and lenticular nuclei – first order thalamic nuclei (that receive driver afferent connections from peripheral sensory receptors but not from cerebral cortex) and pallidum (that does not receive direct anatomical connections from cerebral cortex) exhibit lower degrees of IHFS when compared to their adjacent structures (higher order thalamic nuclei that receive driver afferent connections from cerebral cortex, and putamen that receives direct anatomical connections from cerebral cortex), as shown in Figures 1, 2, 3, Supplementary Figure 2. **(B)** In subcortical structures that lack variability in terms of either direct vs. indirect or driver vs. modulatory cerebral cortical afferent connections, IHFaS is present to a greater degree in regions linked to cerebral cortical territories with higher IHFaS. This conclusion is based on our results in cerebellar cortex and caudate nuclei (Figures 4, 5). Note that the concepts of IHFS/IHFaS are broad, including the fact that two homotopic left and right brain areas may be functionally linked to similar or distinct regions within each structure (as captured by functional gradients that were calculated within each subcortical structure), the related phenomenon that two homotopic left and right brain areas might be functionally linked preferentially to one hemisphere (as captured by laterality indices calculated within each subcortical structure), and the closely linked fact that two homotopic left and right brain areas may be strongly or weakly connected to each other.

### In cerebellar cortex and caudate nuclei, but not in thalami or lenticular nuclei, IHFaS is stronger in areas with connectivity to cerebral cortical areas that also have stronger IHFaS

Analyses of non-principal functional gradients (**Figure 4A**) revealed asymmetric functional gradients in cerebellar cortex and caudate nuclei (**Figure 4B**). These asymmetric functional gradients were related to laterality indices in a manner similar to the thalami and caudate nuclei, namely, more extreme values in asymmetric functional gradients corresponded to territories with stronger IHFaS as indexed by laterality indices (**Figure 4D**). These relationships are observable but not as apparent as in cerebellar cortex and caudate nuclei (**Figure 2**), perhaps due to the fact that asymmetric gradients in cerebellar cortex and caudate nuclei do not correspond to the principal functional gradients (but rather to gradient 3 in cerebellar cortex and gradient 2 in caudate nuclei). In this way, functional connectivity asymmetries in cerebellar cortex and caudate nuclei may not be prominent enough to dictate the direction of the principal axis of macroscale functional variation within these structures.

The observation of weaker IHFS in some territories of cerebellar cortex and caudate nuclei that were identified by asymmetric functional gradients offered an opportunity to interrogate the anatomical and physiological basis of IHFaS in these regions. As noted above, within the lenticular nucleus there are direct cerebral cortical connections to putamen but not to pallidum, and there are driver afferent cerebral cortical connections to higher order but not to first order nuclei within thalamus. In contrast, it is likely that all aspects of caudate nucleus receive connections from cerebral cortex, and that these connections are direct (monosynaptic) (Haber 2016). Similarly, it is likely that all aspects of cerebellar cortex receive and send cerebral cortical projections, and that all these projections possess a uniform circuit architecture, namely, cerebral cortical – ponto – cerebellar cortical afferents, and cerebellar cortical – cerebellar nuclear – thalamic – cerebral cortical efferents (Kelly & Strick 2003; Middleton & Strick 1994; Schmahmann 1996; Schmahmann & Pandya 1991b, 1997b). There also appears to be no evidence of driver vs. modulatory cerebral cortical afferents in caudate or cerebellum, including a lack of variability in the laminar origin, apart from minor variation in sublaminar origin in layer V of cerebral cortical afferents within these structures (corticopontine projections are derived heavily from layer Vb (Glickstein et al. 1985), corticostriatal projections from layer Va more than Vb (Yeterian & Pandya 1994)). It can thus be concluded that there is a contrast between variations in terms of either direct vs. indirect or driver vs. modulatory cerebral cortical afferent connections in thalamus and lenticular nucleus, and an absence of such variations in caudate and cerebellar cortex. This observation raised the possibility that a factor different from properties of cerebral cortical anatomical connections might be associated with IHFS/IHFaS variability in cerebellar cortex and caudate.

Although there are no differences in terms of either direct vs. indirect or driver vs. modulatory cerebral cortical afferent connections within cerebellar cortex and caudate nucleus, there are nevertheless topographically arranged maps of anatomical connectivity linking the cerebral cortex with the cerebellar cortex (Buckner et al. 2011; Guell et al. 2018b; Schmahmann & Pandya 1997a) and caudate nucleus (Parent 1990; Yeterian and Van Hoesen, 1978; Schmahmann and Pandya, 2006; Choi et al. 2012). We interrogated this variable and its relationship to subcortical IHFS/IHFaS by projecting subcortical asymmetric functional gradients to cerebral cortex. This analysis revealed that, in both cerebellar cortex and caudate nuclei, territories of stronger IHFaS are functionally linked to cortical areas with higher IHFaS (predominantly default-mode and frontoparietal networks, **Figure 5**). This observation in cerebellum is consistent with recent reports of stronger functional lateralization in cerebellar territories that correspond to default-mode network (see “symbolic communication” component in cerebellum in Karolis et al. 2019, that corresponds to language areas in cerebellum that also overlap with default-mode territories). Importantly, this association was not present in thalami or lenticular nuclei – within these structures, asymmetric functional gradients projected to cerebral cortical regions that did not exhibit strong IHFaS (predominantly ventral attention network, **Figure 5**).

### Two distinct principles guiding interhemispheric functional symmetry and asymmetry in the subcortex

Our findings led us to conclude that two distinct principles may guide IHFS/IHFaS in the subcortex. In structures with variability in terms of either direct vs. indirect or driver vs. modulatory cerebral cortical afferent connections (thalami and lenticular nuclei), IHFaS is stronger in regions with no direct or driver cerebral cortical afferent connections (**Figure 6A**). In subcortical structures that lack variability in terms of either direct vs. indirect or driver vs. modulatory cerebral cortical afferent connections (cerebellar cortex and caudate nucleus), IHFaS is more prominent in territories linked to cerebral cortical areas with higher IHFaS (**Figure 6B**). These two principles together support a strong relationship between subcortical IHFS/IHFaS and connectivity between subcortex and cortex.

### Future studies

The characterization of subcortical IHFS/IHFaS patterns presented here introduces new questions to basic and clinical neuroscience. An intriguing testable possibility is that IHFS might originate in cerebral cortex and propagate to subcortex. There is no equivalent of the corpus callosum in the subcortex – whereas some fibers link paramedian homotopic subcortical areas, the subcortex is largely devoid of commissural pathways in sharp contrast with the cerebral cortex. IHFS may thus be generated in the cerebral cortex as a result of callosal connectivity and other commissural cerebral cortical fibers (see introduction), and propagated from cerebral cortical homotopic pairs (e.g. left and right M1) to subcortical homotopic pairs (e.g. left and right cerebellar motor areas) through the anatomical connections that link the cerebral cortex with the subcortex. This hypothesis is consistent with our observations (**Figure 6A** and also **Figure 6B**), but the evidence provided here is only correlational. Brain physiology manipulation experiments might provide definitive causal evidence. Our testable hypothesis predicts that alteration of IHFS in cerebral cortex should result in a matching alteration of IHFS in the subcortex, and the same manipulation in subcortex should not change IHFS in the cerebral cortex. For example, increasing IHFS between left and right cerebral cortical M1 should increase IHFS between left and right cerebellar motor cortex, but increasing IHFS between left and right cerebellar motor cortex should result in no or lower increase of IHFS between left and right cerebral cortical M1.

Asymmetry in functional gradients in cerebellar cortex and caudate nucleus was not identified in the first, but rather in the third and second functional gradients, respectively (**Figure 4A**). In contrast, the principal (first) functional gradients in thalami and lenticular nuclei were asymmetric. Differences in IHFS thus appear to dominate the principal axis of functional connectivity pattern variations within thalami and lenticular nuclei, but not within cerebellar cortex and caudate nuclei. Future investigations might explore the reason for this difference. One possibility is that differences in IHFS are more pronounced in structures with more variability in terms of either direct vs. indirect or driver vs. modulatory cerebral cortical afferent connections, and therefore more pronounced in thalami and lenticular nuclei when compared to cerebellar cortex and caudate nuclei. A testable hypothesis is that the principle presented in **Figure 6A** dictates subcortical IHFS variability to a stronger degree than the principle presented in **Figure 6B**. Manipulation studies may provide definitive causal evidence to support this hypothesis: disruption of anatomical tracts linking cerebral cortex to subcortex (**Figure 6A**) should disrupt subcortical IHFS more than physiological disruption of cerebral cortical IHFS (**Figure 6B**). For example, lesion of fibers linking cerebral cortical M1 to cerebellar motor cortex should disrupt cerebellar IHFS more than a functional manipulation of IHFS between left and right cerebral cortical M1.

Future studies may examine whether inter-subject differences exist in the observations reported here and explore the significance of this aspect of brain inter-subject variability in heath and disease. Possible correlates of inter-subject variations in our observations may include presence versus absence of commissural tracts (e.g. interthalamic adhesion is absent in a fraction of the healthy population; Damle et al. 2017), structural properties of commissural fibers (e.g. diffusion-weighted MRI signal in caudate relates to task-based functional asymmetry in cerebral cortex; Karolis et al. 2019), or differences in functional organization that relate to functional lateralization (e.g. there are emotion processing lateralization differences between male and female healthy participants; Canli et al. 2002). Subcortical IHFS inter-subject variability might also be a relevant brain correlate of neurological or psychiatric disorders; previous studies have described homotopic synchrony differences in cocaine addiction (Kelly et al. 2011; Wang et al. 2019). It is also possible that these differences relate to other disease-specific brain correlates such as subcortical volumetric asymmetries (e.g. pallidal volume asymmetries in schizophrenia Okada et al. 2016). Variations in IHFS across individuals might also relate to prognosis of response to therapeutic interventions; recent evidence indicates that integrity of anatomical interhemispheric connections is associated with better response to ipsilateral cerebral cortical stimulation in the treatment of neglect after stroke (Nyffeler et al. 2019).

Our findings indicate that there is lower IHFS in first order compared to higher order thalamic nuclei. Previous studies have described other physiological properties that are different between first and higher order thalamic nuclei, including distinctive single-neuron electrophysiological firing in first order nuclei (Ramcharan et al. 2005). In addition, there are aspects of variability in thalamic cerebral cortical connections other than the distinction between first and higher order nuclei. Tract-tracing experiments in rat reveal that some thalamic nuclei receive projections exclusively from ipsilateral cortical areas, while others receive projections from both ipsilateral and homotopic contralateral cerebral cortical structures (Mathiasen et al. 2017). Future studies might examine the relationship between these distinct properties in first versus higher order thalamic nuclei, as well as the links between IHFS variability and variability in symmetric versus asymmetric cerebral cortical anatomical connectivity in thalamus.

## Conclusion

IHFS is an important property of brain organization with implications for both basic and clinical neuroscience. There is a scarcity of studies describing IHFS in the subcortex, and the proposed anatomical basis of interhemispheric symmetry in the cerebral cortex that is based on commissural connectivity is incompatible with subcortical anatomy. Here we present a detailed description and analysis of IHFS/IHFaS in thalamus, lenticular nucleus, cerebellar cortex, and caudate nucleus. Our observations are encapsulated in the following two principles: (i) in subcortical structures with variability in terms of either direct vs. indirect or driver vs. modulatory cerebral cortical afferent connections (thalami and lenticular nuclei), IHFaS is stronger in regions with no direct or driver cerebral cortical afferent connections; (ii) in subcortical structures with no variability in terms of either direct vs. indirect or driver vs. modulatory cerebral cortical afferent connections (cerebellar cortex and caudate nucleus), IHFaS is more prominent in territories linked to cerebral cortical areas that have higher IHFaS. These observations prompt new questions in basic and clinical neuroscience, and generate novel testable hypotheses for future study.

---

**Supplementary Figure 1.**
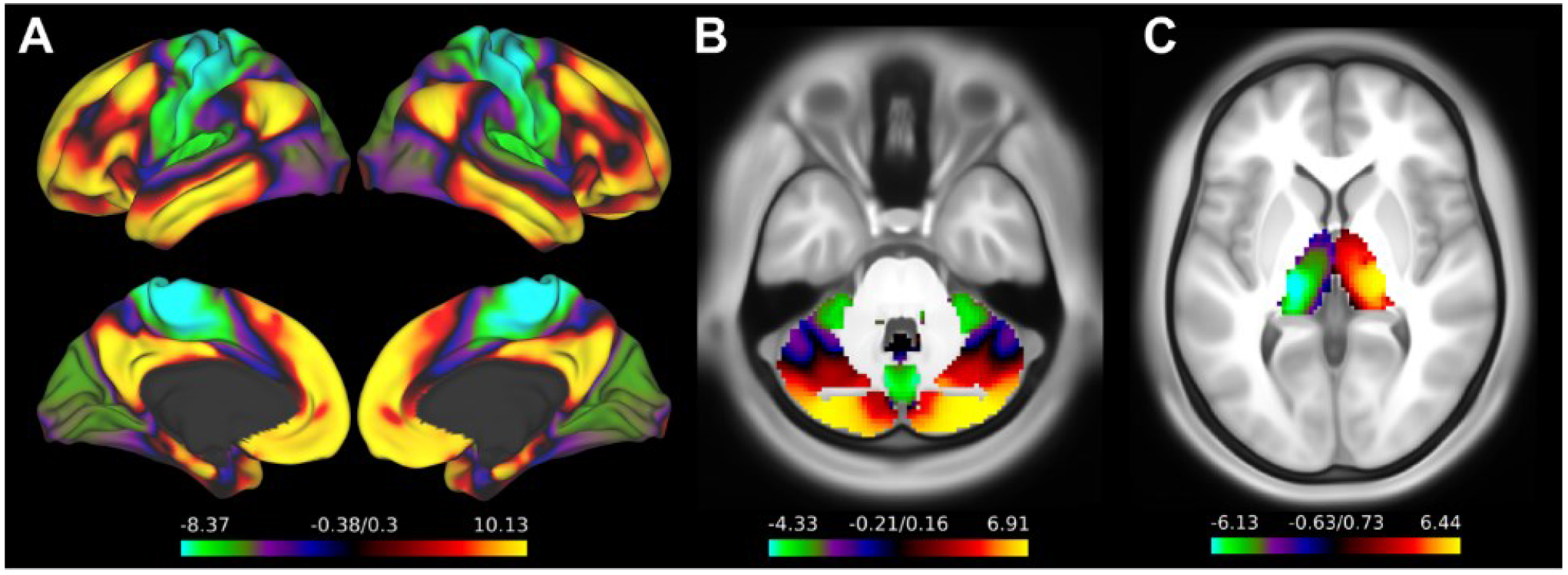
Principal functional gradients are symmetric in cerebral cortex (**A**) and cerebellar cortex (**B**), but strikingly asymmetric in thalamus (**C**; note distribution from left thalamus [green/blue colors] to right thalamus [red/yellow colors]). Cerebral and cerebellar cortical gradients were calculated as reported in (Margulies et al. 2016) and (Guell et al. 2018b), respectively. Thalamic gradients were calculated as described in the methods section.

**Supplementary Figure 2.**
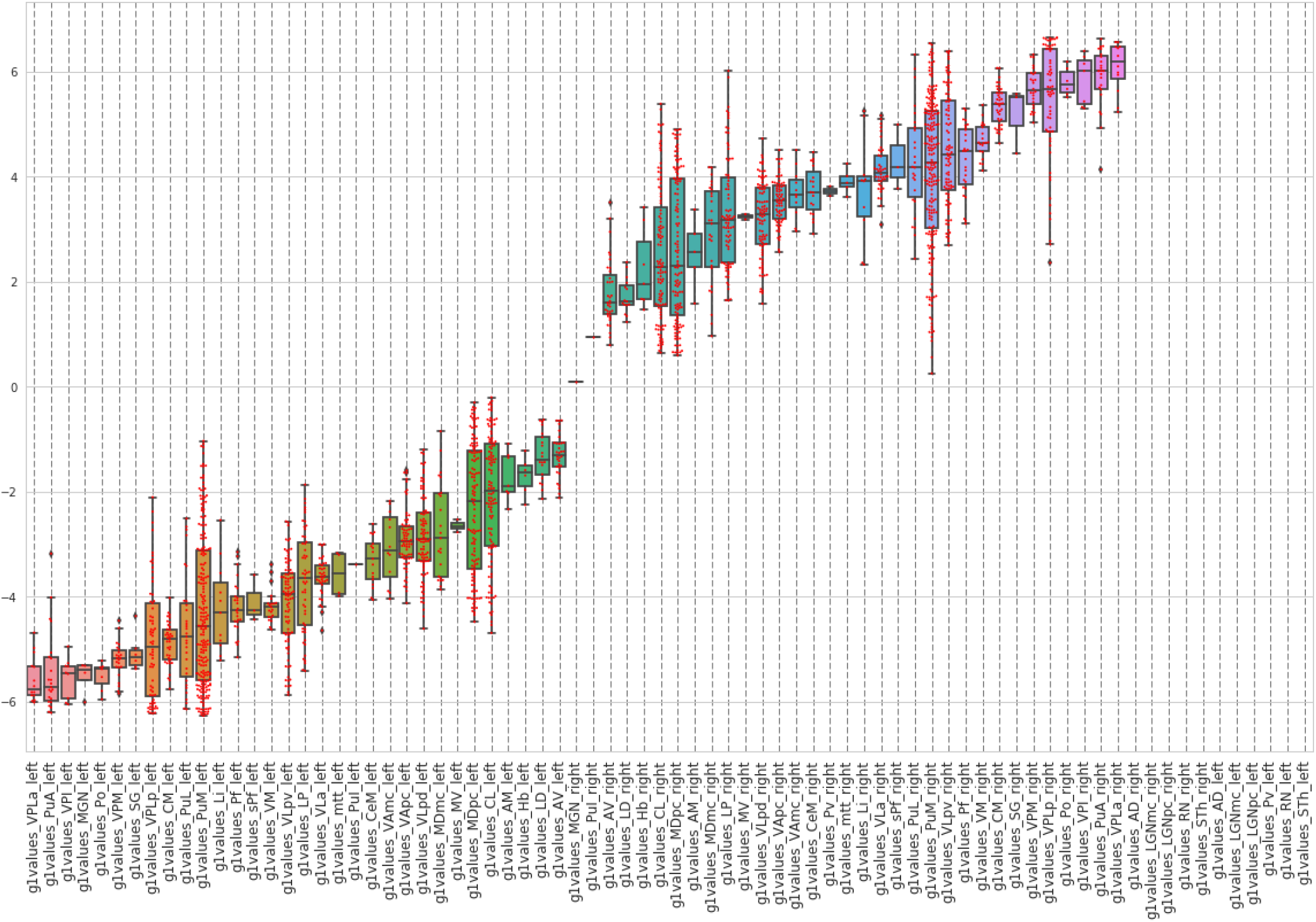
<u>Thalamic principal functional gradient values for each thalamic nucleus</u>. Each red dot corresponds to one voxel in left or right thalamus. Some nuclei were too small and did not obtain a functional gradient value when the MRI anatomical atlas of thalamic nuclei was downsampled to match the resolution of HCP resting-state data (thalamic atlas resolution was higher than resting-state data resolution, and for this reason functional gradient values could not be assigned to the smallest thalamic nuclei; these nuclei are shown at the right extreme of this figure, with no datapoints shown in their corresponding columns). Voxels that overlapped with neighboring nuclei after downsampling were also excluded. A perfect progression is observed from left thalamic nuclei (left side of the figure) to right thalamic nuclei (right side of the figure). Extreme (lowest and highest) values correspond mostly to first order thalamic nuclei including ventral posterior nuclei which receive driver afferent fibers from body and head somatosensory tracts (Felten et al. 2016; Guillery 1995); higher order nuclei such as medial dorsal and anterior nuclei which receive driver afferent fibers from cerebral cortex (Felten et al. 2016; Guillery 1995) are located more centrally in this distribution. A detailed description of first order versus higher order thalamic nuclei and driver versus modulatory fibers is provided in the discussion section. Nuclei abbreviations correspond to the terminology in the atlas developed by Krauth and colleagues (Krauth et al. 2010).

## ACKNOWLEDGEMENTS

Carmen Varela for valuable input regarding first order and higher order thalamic nuclei.

## Notes

### Competing Interest Statement

The authors have declared no competing interest.

